# Genomic landscape of oxidative DNA damage and repair reveals regioselective protection from mutagenesis

**DOI:** 10.1101/168153

**Authors:** Anna R Poetsch, Simon J Boulton, Nicholas M Luscombe

## Abstract

DNA is subject to constant chemical modification and damage, which eventually results in variable mutation rates throughout the genome. Although detailed molecular mechanisms of DNA damage and repair are well-understood, damage impact and execution of repair across a genome remains poorly defined. To bridge the gap between our understanding of DNA repair and mutation distributions we developed a novel method, AP-seq, capable of mapping apurinic sites and 8-oxo-7,8-dihydroguanine bases at ∼300bp resolution on a genome-wide scale. We directly demonstrate that the accumulation rate of oxidative damage varies widely across the genome, with hot spots acquiring many times more damage than cold spots. Unlike SNVs in cancers, damage burden correlates with marks for open chromatin notably H3K9ac and H3K4me2. Oxidative damage is also highly enriched in transposable elements and other repetitive sequences. In contrast, we observe decreased damage at promoters, exons and termination sites, but not introns, in a seemingly transcription-independent manner. Leveraging cancer genomic data, we also find locally reduced SNV rates in promoters, genes and other functional elements. Taken together, our study reveals that oxidative DNA damage accumulation and repair differ strongly across the genome, but culminate in a previously unappreciated mechanism that safeguards the regulatory sequences and the coding regions of genes from mutations.

## 1. Introduction

The integrity of DNA is constantly challenged by damaging agents and chemical modifications. Base oxidation is a frequent insult that can arise from endogenous metabolic processes as well as from exogenous sources such as ionizing radiation. At background levels, a human cell is estimated to undergo 100 to 500 such modifications per day, most commonly resulting in 8-oxo-7,8-dihydroguanine (8OxoG) ^1^. At steady state, up to 2,400 8OxoG sites per cell are reported ^2^. However, estimates differ widely due to differences in methodology ^3,4^. Left unrepaired, 8OxoG can compromise transcription ^5,6,7^, DNA replication ^8^, and telomere maintenance ^9^. Moreover, damaged sites provide direct and indirect routes to C-to-A mutagenesis ^10^.

Oxidative damage is reversed in a two-step process through the base excision repair (BER) pathway ^11^. The damaged base is first recognized and excised by 8-oxoguanine DNA glycosylase 1 (OGG1), leaving an apurinic site (AP-site). Glycohydrolysis is highly efficient, with a half-life of 11 min ^12^. AP-sites are removed through backbone incision by AP-lyase (APEX1), end processing through flap-endonuclease 1 (FEN1), and the base is subsequently replaced with an undamaged nucleotide. Alternatively, in short-patch base excision repair, the base is replaced dependent on polymerase beta. Other sources of non-radiation-induced AP-sites include spontaneous depurination and excision of non-oxidative base modifications, such as uracil. Cells are reported to typically present with a steady state of ∼15000 to ∼30000 AP-sites per cell, which includes the associated beta-elimination product ^2,13^. In response to ionizing radiation, oxidative damage levels increase by several orders of magnitude ^14,15^.

Though originally controversial ^16,17^, there is now broad acceptance that mutation rates vary across different regions within genomes. Background mutation rates in Escherichia coli genomes were shown to vary non-randoμly between genes by an order of magnitude, with highly expressed genes displaying lower mutation rates ^18^. In cancer genomes, single nucleotide variants (SNVs) tend to accumulate preferentially in heterochromatin ^19,20^. More recently, it was reported that SNV densities in cancers are lower in regions surrounding transcription-factor-binding, but are elevated at the binding sites themselves and at sites with a high nucleosome occupancy ^21–24^. Although these variabilities remain mechanistically unexplained, they likely arise through a combination of regional differences in damage sensitivity and the accessibility to the DNA repair machinery ^25^. However, since mutations represent the endpoint of mutagenesis, it is impossible to tease apart the contributions from damage and repair using re-sequencing data alone.

The role of oxidative damage in regional differences of mutagenesis remains largely unclear. Antibodies have been used to study the genome wide distribution of 8OxoG. While the specificity of this approach remains questionable ^26,27^, 8OxoG accumulation was found in GC and CpG island rich, early replicating DNA ^28^, but then also in gene deserts and the nuclear periphery ^29^. Under conditions of hypoxia, 8OxoG accumulates in activated promoters linked to specific transcription factors ^30^. These apparently contradicting conclusions may be explained through different levels of resolution and the fast turnover of 8OxoG into the more persistent AP-sites, oxidative damage that is hidden from 8OxoG detection.

To further our understanding of the molecular mechanisms underlying local heterogeneity of mutation rates, direct and specific measurement of DNA-damage types and intermediates is required at high resolution and on a genomic scale. Dissecting these mechanisms will help understand the local sensitivities of the genome and why certain regions appear to be protected.

## 2. A genome-wide map of oxidative damage

To measure oxidative damage and its intermediates across the genome, we developed an approach that specifically detects AP-sites using a biotin-labelled aldehyde-reactive probe under pH neutral conditions ^13,31^; (Figure 1A, Supplementary Figure S1, and Supplementary Figure S2). Besides the use to detect AP-sites, the same probe has previously been used to measure 5-formyl-cytosine (5-fC), changing the reactivity with an acidic environment ^32^. Raiber et al demonstrated that 5-fC is generated primarily in CpG islands at early development during which the genome is demethylated ^33^. This work confirmed previous studies ^13,31^ that the probe is highly specific for 5-fC at pH5, whereas at pH7 the specificity shifts to AP-sites, which is the experimental condition we use (see SupFig 2 in Raiber et al ^32^); therefore, 5-fC is not expected to be a major contributing factor to measurements in the current study. After fragmentation of genomic DNA, biotin-tagged DNA with the original damage sites was pulled down using streptavidin magnetic beads and prepared for high-throughput sequencing. The signal was quantified as the Relative Enrichment of the pull-down over the input DNA, with positive values indicating regions of damage accumulation. Broad distribution of damage and the gradual changes over the genome does not suggest peaks beyond hot spots in repetitive elements (see below and Figures 1D, 3G, 3H). Therefore a binning approach was chosen, added to by an analysis strategy that addresses damage distribution relative to genomic features, which is analogous to similar studies such as research addressing the specificity of the DNA methylation machinery ^34^.

**Figure 1.**
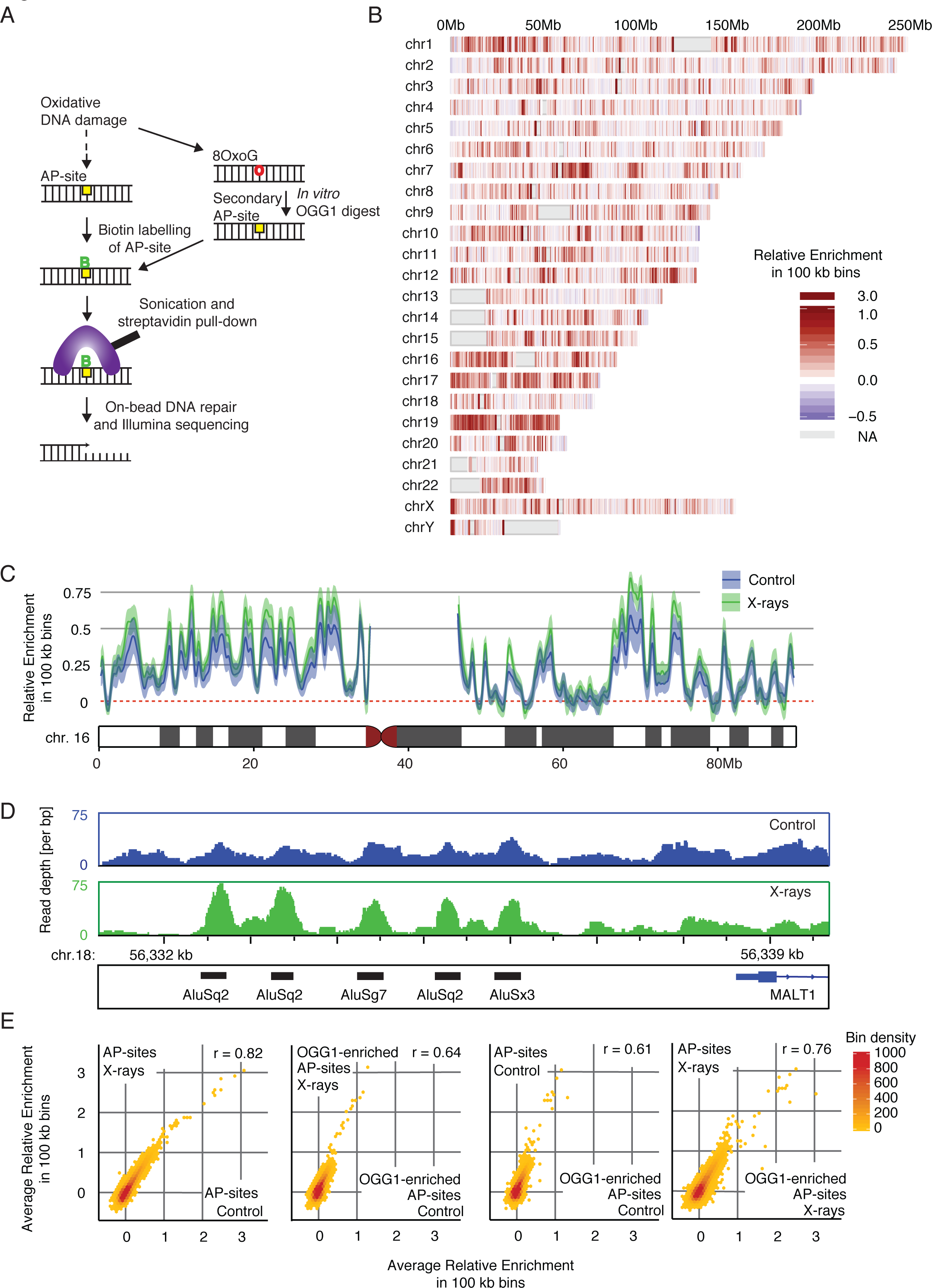
Oxidative damage is heterogeneously distributed at different scales of resolution. (**A**) Schematic of AP-seq, a new protocol to detect apurinic-sites (AP-sites) as a measure of oxidative damage in a genome. 8OxoG is excised by OGG1 in the first, rapid step of base excision repair, leaving an AP-site. DNA containing these sites are biotin-tagged using an aldehyde reactive probe (ARP), fragmented, and pulled-down with streptavidin. The enriched DNA is processed for sequencing and mapped to the reference genome. The damage level across the genome is quantified by assessing the number of mapped reads. To check for unprocessed 8OxoG, we perform an *in vitro* digest of extracted genomic DNA with OGG1 and repeat the AP-site pull-down. **(B)** Genome-wide map of AP-site distribution after X-ray treatment. The colour scale represents the Relative Enrichment of AP-sites in 100kb bins across the human genome, averaged across biological replicates. Grey regions represent undefined sequences in the human genome, such as centromeres and telomeres. Damage levels are highly correlated between treatment conditions at 100kb resolution. **(C)** More detailed view of AP-site distribution on Chromosome 16. Plot lines depict the average Relative Enrichment for X-ray treated (green) and untreated (blue) samples. Shaded boundaries show standard error of the mean for the biological replicates. Untreated and X-ray treated samples display very similar damage profiles. (**D**) Genome browser views of damage distributions for untreated and X-ray treated samples across an 8kb region upstream of MALT1. Damage levels are represented as unnormalised sequencing depth of the pooled biological replicates. At high resolution, it becomes apparent how sharp the damage levels rise over background at *Alu* elements after X-ray treatment, which leads to more distinct patterns than the broader distributed untreated control. (**E**) Scatterplots of the correlation in average Relative Enrichments of samples with differing treatment and OGG1-enrichment conditions. Damage levels are highly correlated across all conditions.

**Figure 2.**
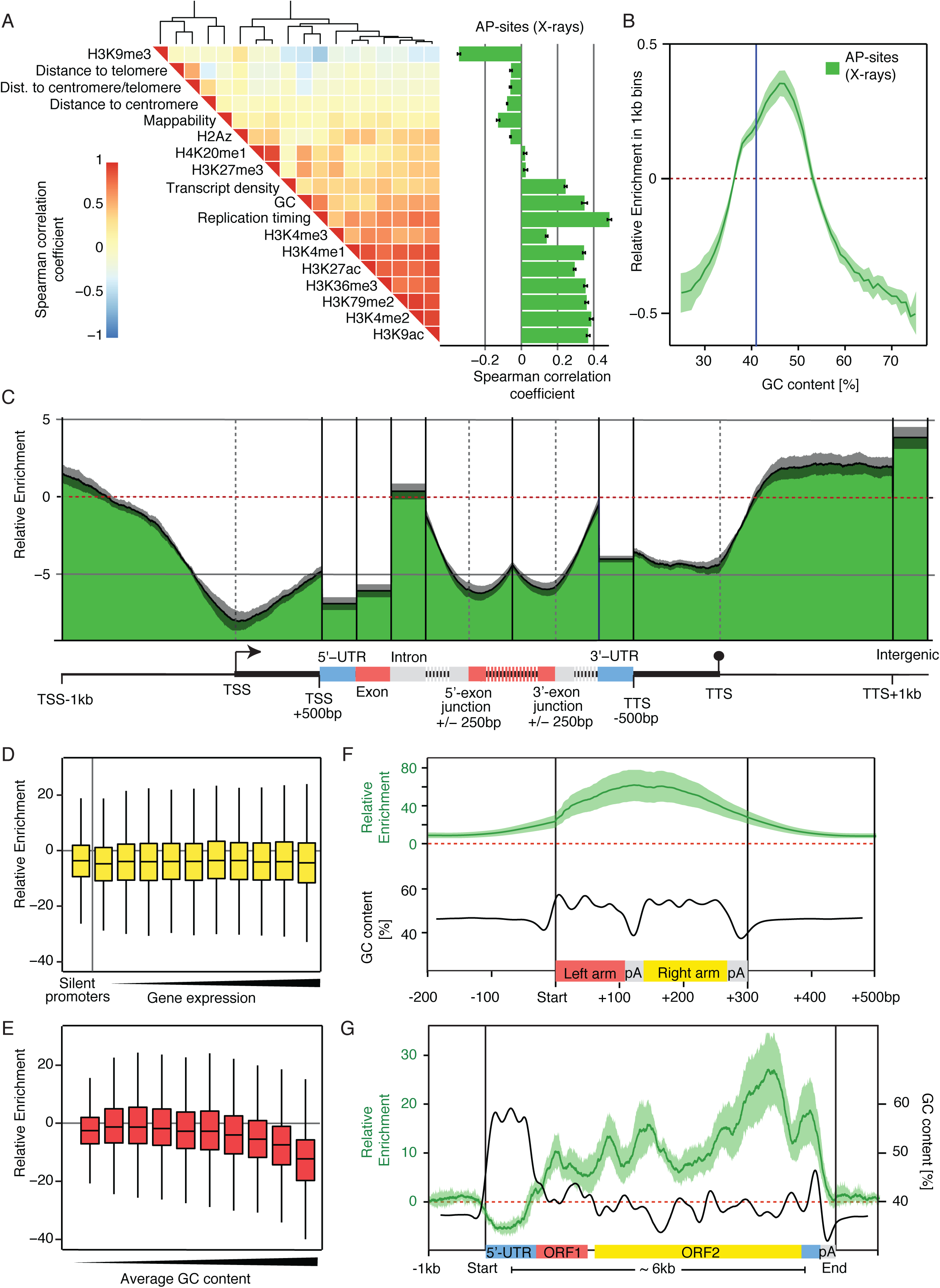
Oxidative damage distribution is associated with genomic features. (**A**) Bar plot displays the average correlation of damage levels with large-scale chromatin and other features in HepG2 cells at 100kb resolution. Damage correlates with euchromatic features and anticorrelates with heterochromatic ones, the opposite of that observed for cancer SNVs. The heatmap shows the relationship between the features, grouped using hierarchical clustering. **(B)** The plot shows dependence between Relative Enrichment of damage and genomic GC content at 1kb resolution. Damage levels increase with GC content and then surprisingly fall in high GC areas. The blue line marks the genomic average GC content of 41%. **(C)** Metaprofile of Relative Enrichment over ∼23,000 protein-coding genes (n_genes_=23,056, n_promoters_=48,838, n_5UTRs_=58,073, n_exons_=214,919, n_introns_=182,010, n_3UTRs_=28,590, n_termination_=43,736, n_intergenic_=22,480). Damagelevels for UTRs, exons, introns, and intergenic regions are averaged across each feature due to their variable sizes. Coding and regulatory regions are depleted for damage, whereas introns have near intergenic damage levels. (**D, E**) Boxplots depict damage levels at 48,838 promoters binned into unexpressed and expression deciles (D), and average GC content deciles (E). Promoters are defined as the transcriptional start sites +/-1kb. Damage is not transcription-dependent, but reduces with increasing promoter GC content. (F, G) Metaprofiles of Relative Enrichments and average GC contents across 848,350 *Alu* and 2,533 *LINE* elements. There is a very large accumulation of damage inside these features. All panels display measurements for X-ray treated samples. Error bars and shaded borders show the standard error of mean across biological replicates.

Figure 1B provides the first high-resolution, genome-wide view of AP-sites following X-ray induced oxidative damage. Increase in damage levels have been confirmed using colourimetric measurements for AP-sites (Supplementary Figure S2A) and immunostaining for gH2AX foci (Supplementary Figure S2B). Measurements of AP-sites represent both the background levels in the genome and those acquired in response to X-ray treatment in HepG2 cells with good reproducibility (Supplementary Figure S3). It immediately highlights the extreme variability in the relative density of AP-sites across the human genome: though the genome-wide mean Relative Enrichment is 0.1, local enrichments vary from less than −0.6 to more than 3.0. Hot and cold spots are found across all chromosomes and do not appear to follow a particular distribution pattern: whereas chromosome 19 presents damage hot spots throughout the chromosome, on chromosome 7 we observe pericentromeric hot spots. Figure 1C shows a more detailed profile of chromosome 16, including distributions for treated and untreated samples. The profiles of the X-ray treated samples indicate an overall treatment-dependent accumulation of damage; however local relative distribution patterns of preexisting background damage are maintained, suggesting that hot spots gain the most additional damage. In Figure 1D, we zoom further into an 8kb region upstream of the MALT1 gene. Here, differences between the treated and untreated samples become apparent, with damage after X-ray exposure particularly accumulating on *Alu* transposable elements in comparison to the surrounding sequence. Whereas background AP-site levels indicate a similar trend, the more evenly distributed coverage shows less specific enrichment in *Alu* sequences. These plots exemplify how variable damage enrichments can be, with hot and cold spots occurring from ∼50-500bp to kilo base resolution.

To assess whether the distribution of AP-sites is representative of 8OxoG we applied recombinant OGG1 *in vitro* to the extracted DNA (Figure 1A); under the chosen conditions, any remaining 8OxoG is excised after DNA extraction largely sequence non-specifically ^35^ to result in a set of secondary AP-sites and to a lesser extent the associated beta-elimination product ^36^. In vitro, oligo-nucleotides with 8OxoG derived secondary AP-sites were pulled down with 12.1% recovery rate relative to input, an 11-fold increase as compared to the oligonucleotide containing guanine (Supp. Figure 2). This 1.1% recovery rate represents the technical background level for oligonucleotides, which is to a large part due to heat induced DNA damage, prompted by the oligonucleotide annealing step.

With the conversion of 8OxoG into AP-sites, both damage types are measured simultaneously. However, any difference in enrichment patterns between the original and OGG1-enriched samples indicates the presence of unprocessed 8OxoG in vivo. Although quantitatively different, the control and X-ray treated samples are highly correlated overall (Figure 1E). Moreover, the OGG1-enriched samples are very similar to the primary AP-sites, indicating that at 100kb resolution the AP-site pull-down after irradiation provides a good representation of total oxidative damage patterns. Therefore, the AP-site measurements after X-ray treatment, the sample with the most pronounced patterns, is shown as representative in the following analyses.

## 3. Genomic features shape distribution of oxidative damage

### 3.1 Damage accumulates preferentially in euchromatin but not heterochromatin

To identify potential causes of variation across the genome we compiled for the same HepG2 cell line a set of 18 genomic and epigenomic features associated with DNA damage, repair, and patterns of mutagenesis (Figure 2A). Previous studies reported that SNV densities in cancer genomes were positively correlated with heterochromatic markers (eg, H3K9me3) and negatively correlated with euchromatic ones (eg, H3K4me3, H3K9ac) ^20^. Here, AP-sites display the opposite trend, correlating with open chromatin and anticorrelating with closed chromatin as previously suggested for 8OxoG ^28^. At first glance, it is surprising that SNVs and DNA damage should show opposing trends; However, there are multiple steps of mutagenesis in between, which act chromatin dependently, e.g. repair efficiencies ^37,38^ and replication accuracy ^23^, the impact of which would have to be determined experimentally. Observations are upheld at higher resolutions for many features; for instance, the Spearman’s correlation with H3K9me3 is −0.48 at 1Mb resolution, −0.34 at 100kb, −0.3 at 10kb, and −0.14 at 1kb resolution. For other features, these correlations break down; DNase I hypersensitivity correlates at low resolution (Spearman’s r = 0.5 and 0.3 at 1Mb and 100kb respectively), but the relationship is lost at higher resolutions (r = 0.06 and −0.06 at 10kb and 1kb respectively). This suggests that more detailed genomic features and functional elements also play a role in shaping the local damage distributions.

### 3.2 Damage enrichment is GC-content dependent

As oxidative damage predominantly occurs on guanines ^1^, base content is expected to be a prime determinant of genome-wide distribution. The heatmap in Figure 2A shows that this is true in general, with average damage levels in 100kb windows correlating with GC content (Spearman’s r = 0.37). However closer examination shows a more complex relationship: in Figure 2B, we plot average damage levels in 1kb windows against their GC content. While there is a clear increase in damage as GC content rises from 25% to 47%, this relation breaks down above 47% GC and damage levels drop sharply. This indicates that while there is a larger proportion of the receptive base with increasing GC content, damage in regions of high GC content cannot be explained by base composition alone.

### 3.3 Gene promoters and bodies show selective protection from damage

Next, we interrogated damage distributions over coding regions by compiling a metaprofile for 23,056 protein-coding genes (Figure 2C). The analysis reveals rigid compartmentalisation, with relative damage levels varying substantially between elements. Damage is dramatically reduced within genes compared with flanking intergenic regions (Relative Enrichment = 3.8), most prominently at the transcriptional start (Relative Enrichment = −8.0), 5’-UTRs (Relative Enrichment = −6.9), exons (Relative Enrichment = −6.1) and termination sites (Relative Enrichment = −5.8). In stark contrast, introns show high damage (Relative Enrichment = 0.4), though still below intergenic levels. Intron-exon junctions are accompanied by steep transitions in damage indicating the sharp distinction between coding, regulatory and non-coding regions (Relative Enrichment changes from −6.0 to −0.5 within 300bp around the 3’-exon junction). Damage levels rapidly rise again downstream of termination sites towards intergenic regions (Relative Enrichment shifts from −4.3 to 2.0 within 500bp).

Promoters and transcription start sites have the lowest damage levels of any functional element in the genome (average Relative Enrichment = −8.0 compared with intergenic average of 3.8), similar to what has been shown for alkylation adducts in yeast ^38^. Unlike SNVs and other damage types, which decrease with rising expression levels, we do not detect an association between oxidative damage and expression (Figure 2D). There is a substantial GC content effect (Figure 2E); but in contrast to expectations from base composition alone, damage levels fall as GC content rises (Relative Enrichment = 1.1 at 45% GC and Relative Enrichment = −12.6 at > 64% GC).

### 3.4 Retrotransposons accumulate large amounts of damage

Retrotransposons ^39^ provide a fascinating contrast to coding genes: Long Interspersed Nuclear Elements (LINEs) possess similar structures to genes with an RNA Pol II-dependent promoter and two open reading frames (ORFs), whereas Short Interspersed Nuclear Elements (SINEs) resemble exons in their nucleotide compositions and presence of cryptic splice sites. Unlike coding genes though, LINEs and SINEs accumulate staggeringly high levels of damage. *Alu* elements, the largest family among SINEs, show by far the highest damage levels of any annotated genomic feature: a metaprofile of >800,000 *Alu* elements in Figure 2F peaks at an average Relative Enrichment of 59, much higher than the genomic average of 0.1. The damage profile rises and falls within 500bp. Similarly, a metaprofile of >2,500 *LINE* elements in Figure 2G displays heterogeneous, but high levels of damage accumulation: like coding genes, there is reduced damage at promoters (average minimum Relative Enrichment = −5.2), but in contrast to genes there is a gradual increase in damage from the 5’ to 3’end, peaking at a Relative Enrichment of 26.9 near to the end of the second ORF.

Retrotransposons, though usually silenced through epigenetic mechanisms ^40^, can be activated through loss of repair pathways ^41^ by DNA damage in general ^42^ and ionizing radiation in particular ^43^. How DNA damage or repair affects such silencing mechanisms is currently unknown. One might speculate that DNA damage at these positions could lead to unwanted *LINE* transcription, for instance through repair-associated opening of the chromatin. These distinct and unique damage patterns of both protection and strong accumulation of damage within one functional element suggest the existence of targeted repair or protective mechanisms that are unique to retrotransposons.

### 3.5 Transcription factor-binding sites, G-quadruplexes and other regulatory sites

Finally, we examine the most detailed genomic features previously implicated in mutation rate changes. In Figure 3A-C we assess the impact of DNA-binding proteins: there is a universal U-shaped depletion of damage levels +/-500bp over the binding-site regardless of the protein involved, suggesting that the act of DNA-binding itself is a major protective factor. We find the greatest reduction in damage for actively used binding-sites that overlap with DNase-hypersensitive regions in the HepG2 cell line. However, a smaller reduction is also present for inactive sites, indicating that the effects go beyond simple DNA-binding. It is notable that the binding-site effects override the contribution of the GC content to damage levels.

**Figure 3.**
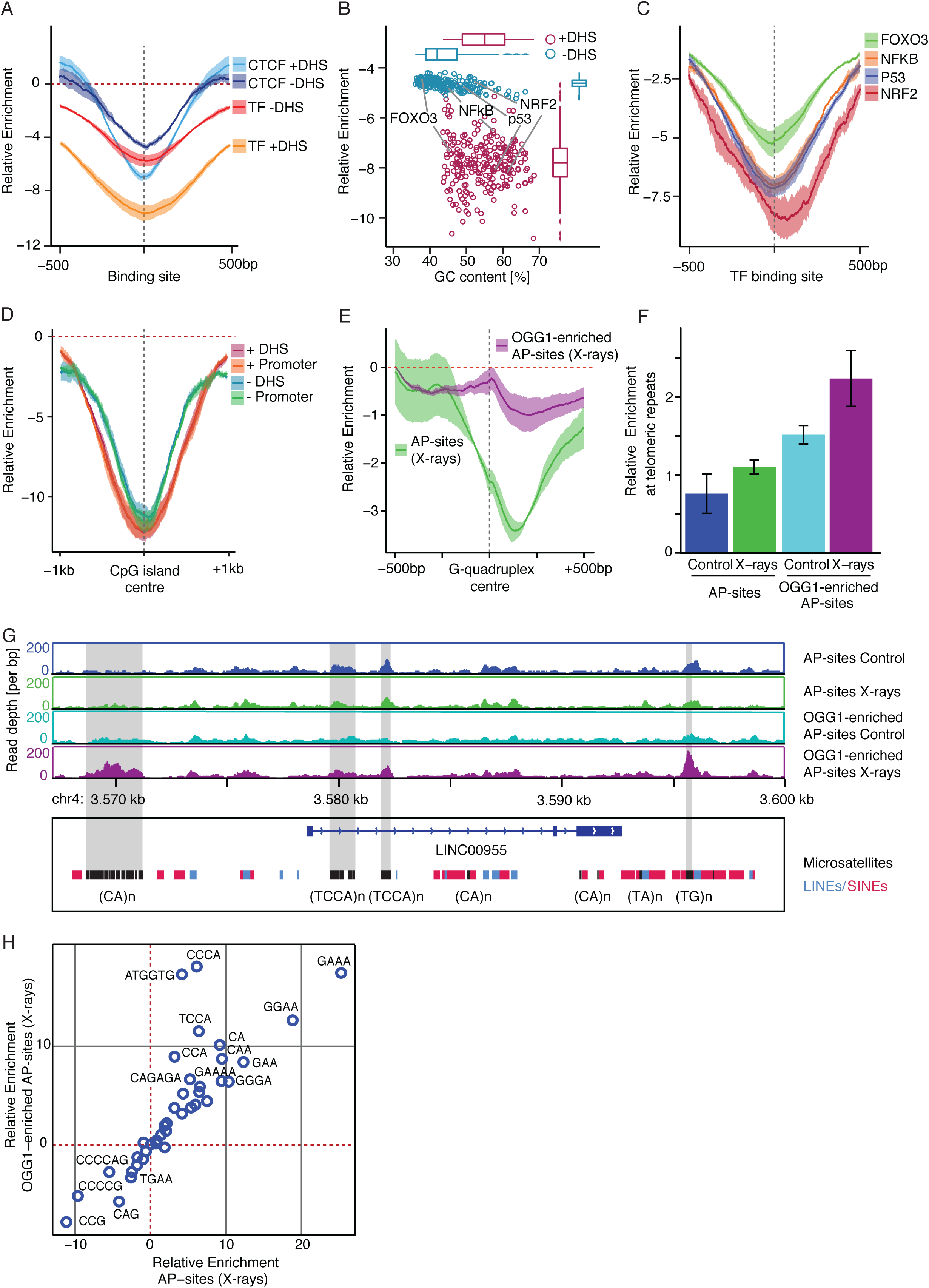
Oxidative damage distribution is associated with regulatory sites and repeats. (**A**) Metaprofiles of Relative Enrichments centered on CTCF-and DNA-binding sites within and outside DNase hypersensitive regions (DHS; n_CTCFinDHS_=37,763, n_CTCFnotDHS_=10,908, n_TFbsInDHS_=253,613, n_TFbsNotDHS_=5,463,612). Damage levels are reduced around binding sites. Shaded borders show the standard error of mean across biological replicates. **(B)** Scatter plot of average Relative Enrichments and GC contents +/-500bp of binding sites for each transcription factor. Binding sites are separated into within and outside DNase hypersensitive sites. Damage levels are universally reduced regardless of transcription factor, with particularly lowered levels for actively used sites in DHS regions. **(C)** Metaprofiles centred on binding sites for 4 selected transcription factors. (**D**) Metaprofiles centred on CpG islands, within and outside promoters and DHS regions (n_DHS_=17,565, n_NotDHS_=9878, n_Promoter_=14850, n_NotPromoter_=12,593). Damage levels are reduced regardless of location and accessibility. (**E**) Metaprofiles centred on predicted G-quadruplexes (n=359,449). There are asymmetrically reduced damage levels for AP-sites, but not for OGG1-enriched AP-sites. (**F**) Bar plots of average Relative Enrichments in G-quadruplexes at telomeric repeats across the 4 treatment and processing conditions. Damage levels are increased in OGG1-enriched samples. Error bars show the standard error of mean across biological replicates. (**G**) Genome browser views of unnormalised damage levels in ∼30kb locus surrounding LINC00955, including microsatellite repeats. Some groups of microsatellites accumulate large amounts of damage and reduced 8OxoG processing. (**H**) Scatter plot displaying average damage levels in different microsatellites types for the AP-site and OGG1-enriched samples. Reverse complementary repeats were assigned to the alphabetically first repeat. Most types display similar damage levels in the two processing conditions; however, several display elevated damage in the OGG1-enriched sample. All panels display measurements for X-ray treated samples, unless indicated otherwise.

GC-rich features are particularly interesting because of the complex relationship between GC content, protein-binding and damage levels. CpG islands are frequently located in promoters and display reduced damage (Figure 3D). Most surprising is the dramatic reduction in damage in CpG islands outside promoters and DNase-hypersensitive regions, indicating that the localisation in promoters is not the main reason for damage reduction; in fact, it is possible that the reduction in damage for high-GC promoters might be explained by the presence of CpG islands and not vice versa.

Another feature of GC-rich sequences are G-quadruplexes (G4 structures) formed by repeated oligo-G stretches. G-quadruplexes are prevalent in promoters ^44^ and telomeric regions ^45^, where they impact telomere replication and maintenance ^46^. A meta-profile for >350,000 predicted G4 structures displays a dramatic asymmetric reduction in damage, in which the minimum occurs just downstream of the G-quadruplex centre (Figure 3E). In line with hypoxia induced 8OxoG accumulation at G4 structures ^30^ we identify G-quadruplexes as one of the few features with clear differences between the 8OxoG and AP-site distributions with a particular enrichment at the centre of G4 structures. This finding is particularly relevant for telomeric repeats (Figure 3F), where oxidized bases impact on telomerase activity and telomere length maintenance ^47^. These repeats are thought to form G4 structures, but in contrast to quadruplexes in general, telomeres present with a mild increase in AP-sites after X-ray treatment (average Relative Enrichment=1.1) and stronger enrichment of OGG1-enriched AP-sites (average Relative Enrichment=2.3).

Micro-satellites are 3-6bp sequences that are typically consecutively repeated 5-50 times. Whereas GC-rich micro-satellite repeats show generally reduced damage, most simple repeats show an accumulation of damage; this is depicted for individual repeat sites at the LINC00955 locus (Figures 3G). The motifs (GAA)_n_, (GGAA)_n_, and (GAAA)_n_ accumulate the largest amounts of damage (Figure 3H). Interestingly, specific sequences display preferential damage enrichment in the OGG1-enriched samples, such as (CCCA)_n_ and (ATGGTG)_n_. Micro-satellites are capable of forming non-B-DNA structures, such as hairpins ^48^; we suggest that changes in the DNA’s local structural properties impairs 8OxoG-processing on these genomic features with possible regulatory functionality.

## 4. SNVs in oxidative damage-dependent cancers reflect underlying damage profiles

Lastly, we address how the distribution of oxidative DNA damage is reflected in the landscape of SNVs in cancer genomic data. We compiled a dataset of 9.4 million C-to-A transversions, the major mutation-type caused by oxidative damage ^49^, from 2,702 cancer genomes ^50^. Of these, 8 hypermutated tumours are defective in polymerase epsilon (Pol E) activity (total 3.4 million C-to-A SNVs). Under normal conditions, Pol E-proofreading prevents 8OxoG-A mismatches, but in the absence of this activity, a large proportion of mismatches are thought to result in C-to-A mutations ^51^. Thus, the distribution of SNVs in the absence of Pol E-proofreading is expected to follow the underlying oxidative damage pattern, reflecting local differences in damage susceptibility and repair preferences ^52^. We also identified 2,401 tumours with increasing proportions of C-to-A SNVs originating from the mutational process associated with the COSMIC Mutational Signature 18, which has been suggested to arise from oxidative damage ^53,54^. 12 tumours harbor coding mutations in enzymes directly involved in 8OxoG or AP-site processing.

In most tumours, about 9% of C-to-A SNVs occur in regions of high GC content (Figure 4A); however, the proportion drops to just 3% among Pol E-defective tumours, in line with the unexpected depletion of oxidative damage in these genomic regions (Figure 2B). Similarly, tumours display decreasing proportions of SNVs with rising amounts of Signature 18 (Figure 4A), following the expected trend for oxidative damage. When mutated in OGG1, APEX1, or FEN1, the proportion rises above average to an average of 11%. We also observed that damage is preferentially distributed in euchromatin at 100kb resolution, whereas SNVs tend to accumulate in heterochromatin; unsurprisingly at this resolution, the damage and SNV densities are anticorrelated (Spearman’s r = −0.49 and −0.45 for proofreading-defective and control tumours respectively).

**Figure 4.**
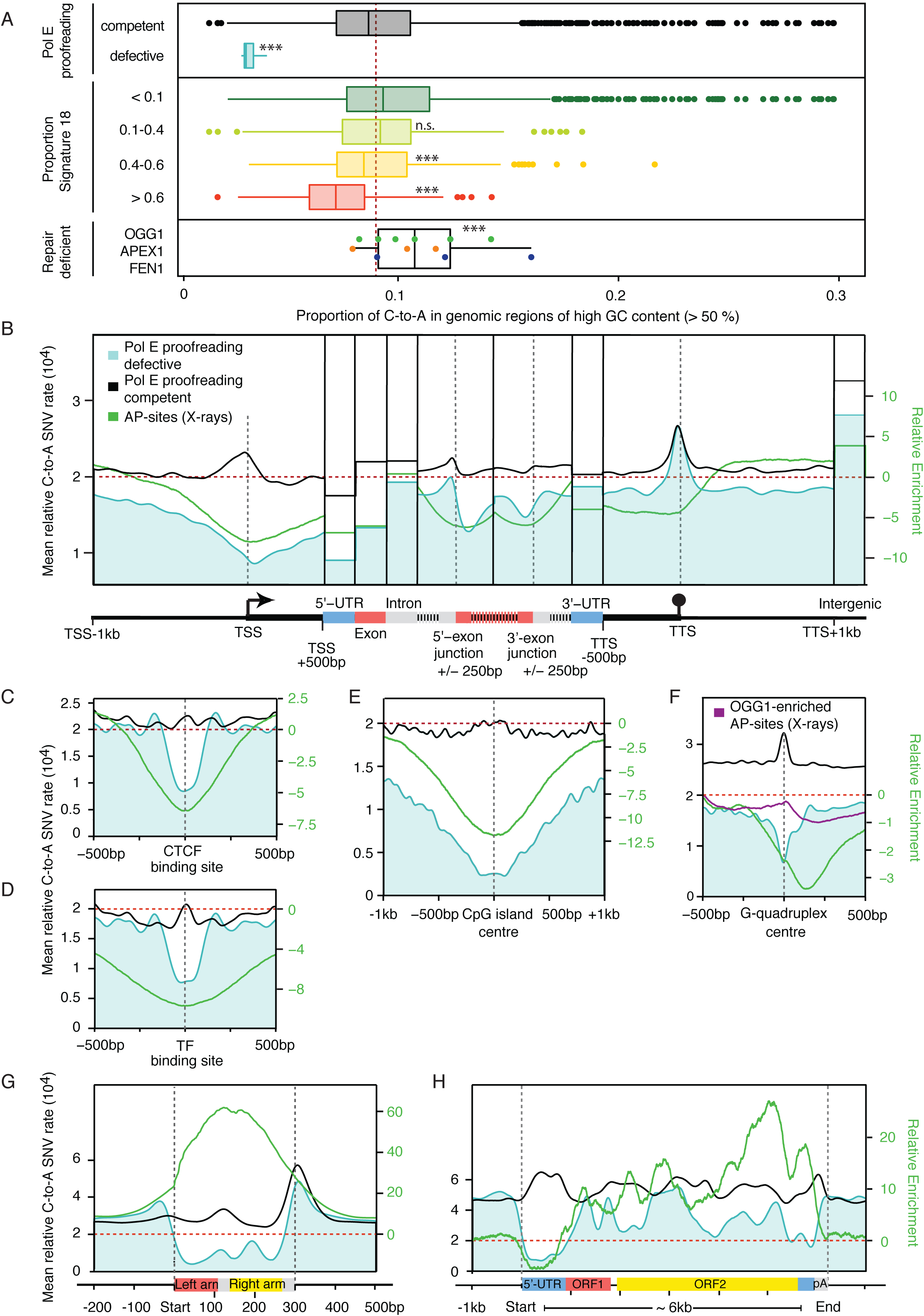
Oxidative damage patterns are reflected in cancer mutagenesis. **(A)** Boxplots of the proportion of C-to-A SNVs (including the reverse complement G-to-T) in genomic regions of high GC content (>50%). Tumour samples are separated into those that are Pol E-proofreading defective (n=8) and to all other tumours (n=2,694), into 4 groups according to Mutational Signature 18 contributions (n_<0.1_=1398, n_0.1-0.4_=322, n_0.4-0.6_=540, n_>0.6_=141), and tumours with coding mutations in OGG1 (n=6), APEX1 (n=3), and FEN1 (n=3). Asterisks indicate significance of p<0.001 by Wilcoxon rank test comparing the PolE proofreading deficient to competent, the different Signature 18 proportions to Signature 18 <0.1, and Repair deficient tumour samples to Signature 18 >0.6. Tumours that are proofreading defective and high in Signature 18 display lower proportions of SNVs in GC-rich regions, while tumours with mutations in OGG1, APEX1, or FEN1 show higher proportions. **(B)** Metaprofile of SNV rates over ∼23,000 protein-coding genes in proofreading defective and control tumours. The damage profile is overlaid for comparison. The oxidative damage-dependent SNV profiles in proofreading-defective tumours show similar distributions to AP-sites, whereas the pattern is lost in control tumours. (**C-F**) Metaprofiles of SNV rates centred on CTCF-binding sites (n=48,671), transcription factor-binding sites in DHS regions (n=253,613), CpG islands (n=27,443), and G-quadruplex structures (n= 359,449). SNV profiles in proofreading defective tumours mimic the damage profiles. (**G, H**) Metaprofiles across 848,350 *Alu* and 2,533 *LINE* elements. SNV rates in proofreading defective tumours are reduced compared with damage profiles.

We focused on the proof-reading defective and control tumour samples for the high-resolution genomic features, as they contain the largest numbers of SNVs. In protein-coding genes, the SNV distribution for Pol E-defective tumours is remarkably similar to the damage profiles (Figure 4B): decreased rates at the TSS, 5’-UTR, exons, and increased rates in introns. The profile is lost in control tumours: we speculate that bulky adducts or strand breaks - a distinct form of damage - cause the accumulation of SNVs at the promoter. SNVs are also depleted from GC-rich genomic features in Pol E-defective tumours, including CTCF-binding sites, transcription factor binding sites, CpG islands and G-quadruplexes. The patterns are lost in the controls (Figure 4C). The difference between the two tumour sets indicates that at high resolution, the distribution of distinct damage types dominates the ultimate SNV profiles. However, there is a striking divergence from damage distributions in retrotransposons (Figure 4G and H); whereas above we observed high levels of damage in Alus and LINEs, there appears to be increased safe-keeping, leading to lower levels of mutations. This pattern is lost in the control tumours.

## 5. Discussion

Our results demonstrate the feasibility of measuring AP-sites across a genome at ∼300bp resolution and high specificity. Damage is strongly reduced in regions of high GC content, which also depends on DNA accessibility. Using the same probe under acidic conditions (pH5) to measure 5-fC, CpG islands have been shown to accumulate this DNA modification in early development ^32^. However, using the probe under neutral pH in HepG2 cells, thus measuring AP-sites, we observe the opposite, a strong reduction of AP-sites in CpG islands. This confirms that the contribution of 5-fC to the measurements through side reactions with the probe under neutral conditions should be negligible.

Previous measurements of oxidative damage using antibodies for8OxoG agree with the accumulation of oxidative damage in open, early replicating DNA ^28^. Other studies describe oxidative damage accumulation in the nuclear periphery and gene deserts ^29^ as well as in certain promoters ^30^. Addressing the more persistend AP-sites, we find open DNA increasingly damaged at the 100kb scale. However, unprocessed 8OxoG accumulates in particular at potential DNA secondary structures, such as G-quatruplexes, telomeres and certain simple repeats. This may explain some discrepancy between AP-site measurements and previous studies on 8OxoG distribution.

In addition to the considerable feature-dependent variability in damage rates, we are able to relate them directly to patterns of SNV occurrences in cancer genomes. At the 100kb scale, euchromatin has increased damage levels, yet fewer SNVs. One could speculate that exposure to oxygen radicals but also better accessibility for repair enzymes, or more accurate replication may lead to this discrepancy, which should be further investigated. At the 10kb to 300bp resolution, we find reduced damage levels in functional elements such as coding sequences, promoters, and transcription factor binding sites, which correlate with SNV occurrences in cancers. The heterogeneity likely results from changes in the balance of damage susceptibility and repair rates at different genomic regions.

Locus-specific oxidative damage is distinct from damage types repaired by other pathways such as nucleotide excision repair (NER). For instance, oxidative damage levels are seemingly independent of gene expression, whereas nucleotide excision repair can be coupled to transcription ^55^. Moreover, for NER, Sabarinathan and Perera reported UV-dependent mutation hotspots around transcription factor binding sites explained by hindered access of the repair machinery. For oxidative damage, we observe the opposite: protection of the same regions from oxidative damage and its derived mutations. Such hotspots are probably prevented through inaccessibility of the DNA to oxygen radicals, which is not the case for UV light. Alternatively, increased repair activity in these regions may lead to a reduction of oxidative damage levels in addition to mismatch repair reducing local mutation rates as described by Frigola ^52^.

Intriguingly, though damage accumulates in LINEs and Alus, they are protected from mutations in cancer genomes; this suggests a specific mechanism for targeted repair at these features that was not reflected in the damage distribution or may be defective in the HepG2 cell line used here. Modulation of DNA damage at these sites would suggest not only effects on mutagenesis, but perhaps even an epigenetic regulatory mechanism through oxidative damage to silence retrotransposons; indeed, an epigenetic function for 8OxoG has been suggested for hypoxia induced gene expression and at G4 structures ^30,56^. At these sites and other potential non-B-DNA structures we detected elevated signals in the OGG1-enriched samples confirming the in vivo accumulation of 8OxoG that had been shown under hypoxic conditions ^30^; this suggests that 8OxoG-processing is impaired. It is interesting to speculate that these sites may have acquired a regulatory function beyond accumulating mutations.

In conclusion, we have established a robust method to measure oxidative damage in a genomewide manner. With minor modifications, it will be suitable for detecting any base modification that can be excised with a specific glycohydrolase. Identifying the pathways that lead to selective repair fidelity and protection of functional elements will not only provide insights into basic mutagenesis but will also allow us to identify any regulatory characteristics of 8OxoG and APsites as epigenetic marks.

## 7. Figures

**Supplementary Figure S1.**
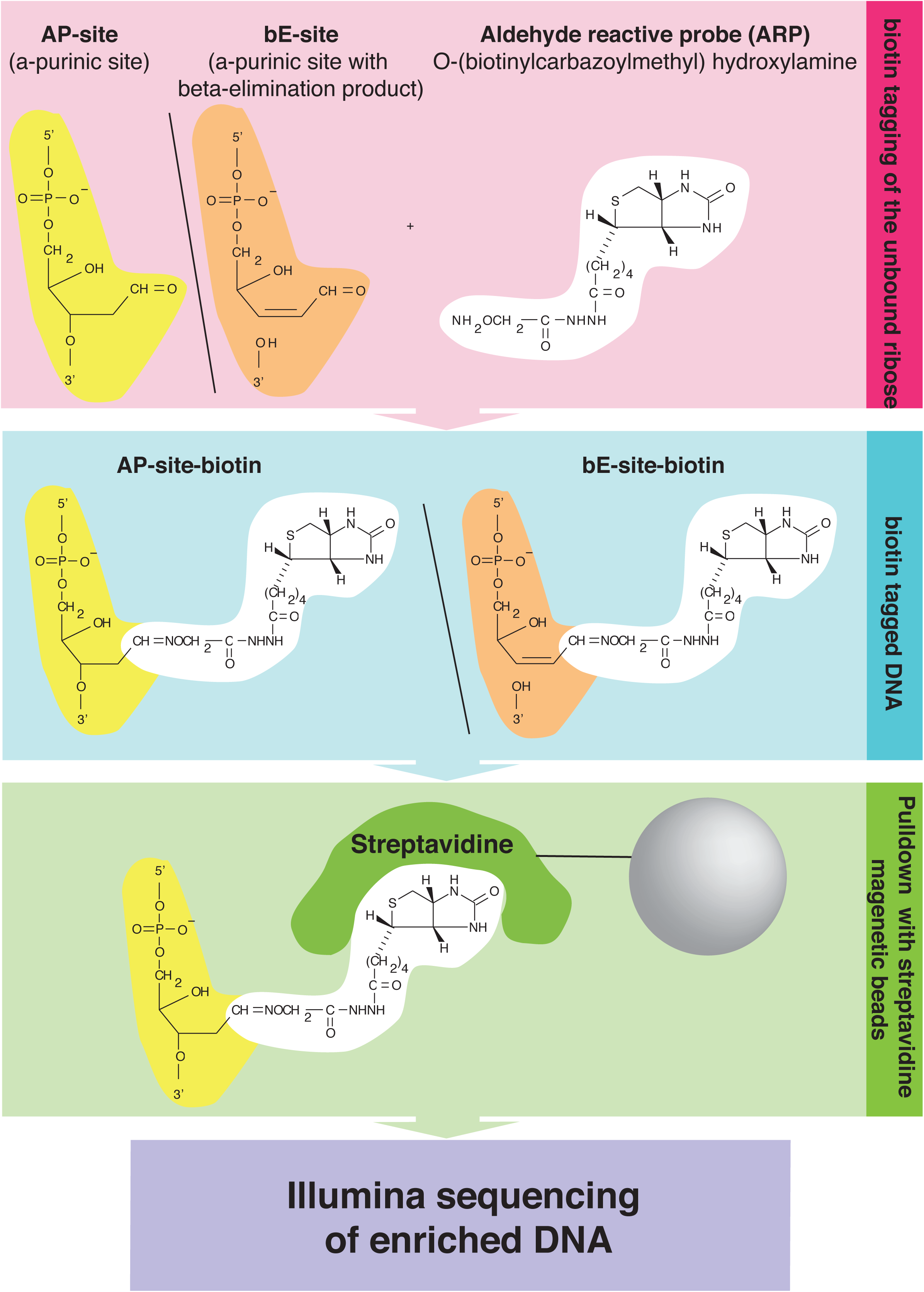
Schematic diagram of the chemical enrichment process of AP-sites using an aldehyde reactive probe. AP-sites and the beta-elimination intermediates of AP-sites are biotin-tagged using an aldehyde reactive probe (ARP) on the damaged strand. Subsequently they are fragmented, and the double stranded DNA is pulled-down with streptavidin. The enriched DNA is processed for sequencing and mapped to the reference genome.

**Supplementary Figure S2.**
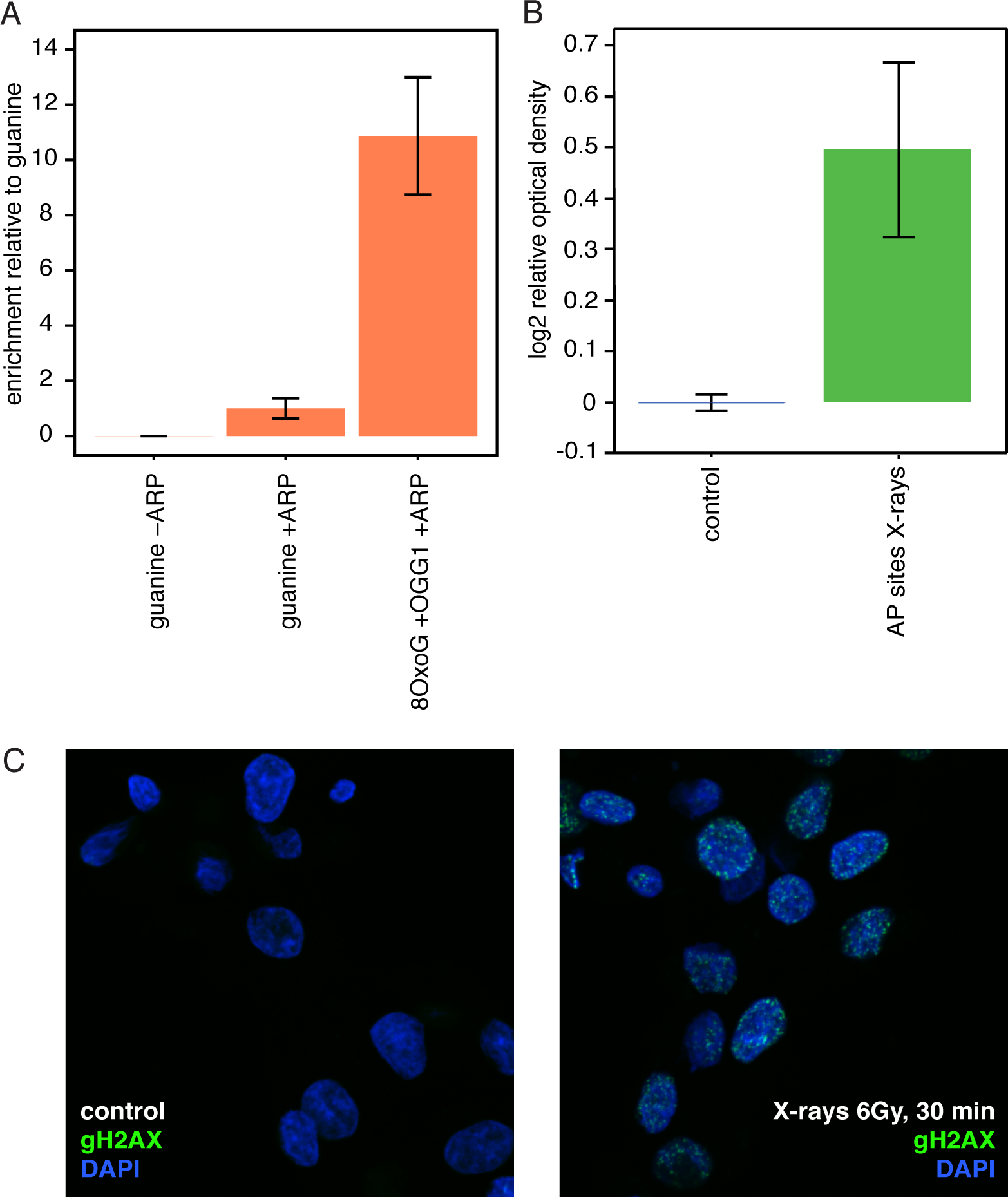
Quality control measures for successful treatment and pulldown specificity. (**A**) In vitro pulldown of standardized oligonucleotides. Oligonucleotides containing 8OxoG converted into an AP-site were pulled down *in vitro* using guanine as control. The DNA was treated in triplicates with OGG1+ARP, ARP alone or was not biotin-tagged. The efficiency of the pulldown was determined as recovery of input and normalised to the guanine control that represents the background levels arising from spontaneous AP-site formation and unspecific probe reaction. Pulldown of untagged DNA is negligible. Pulldown of AP-sites that are created with OGG1 digest of 8OxoG are recovered ∼ 10-fold over undamaged oligonucleotides, which is significant (p < 0.05, student’s t-test). Depicted is the mean and standard error of the mean. **(B)** Colorimetric measurement of AP-sites after X-ray treatment (30 min, 6Gy) using the Aldehyde Reactive Probe. Data are quantified in triplicates as the log2-fold change of normalized optical density relative to the untreated control (p < 0.05, student’s t-test). Depicted is the mean with standard error of the mean. **(C)** Immunofluorescence staining of gH2AX (green) as a measure of radiation induced DNA damage and nuclear staining as a reference (blue) under untreated conditions (left) and after treatment with 6 Gy X-rays and 30 min incubation (right).

**Supplementary Figure S3.**
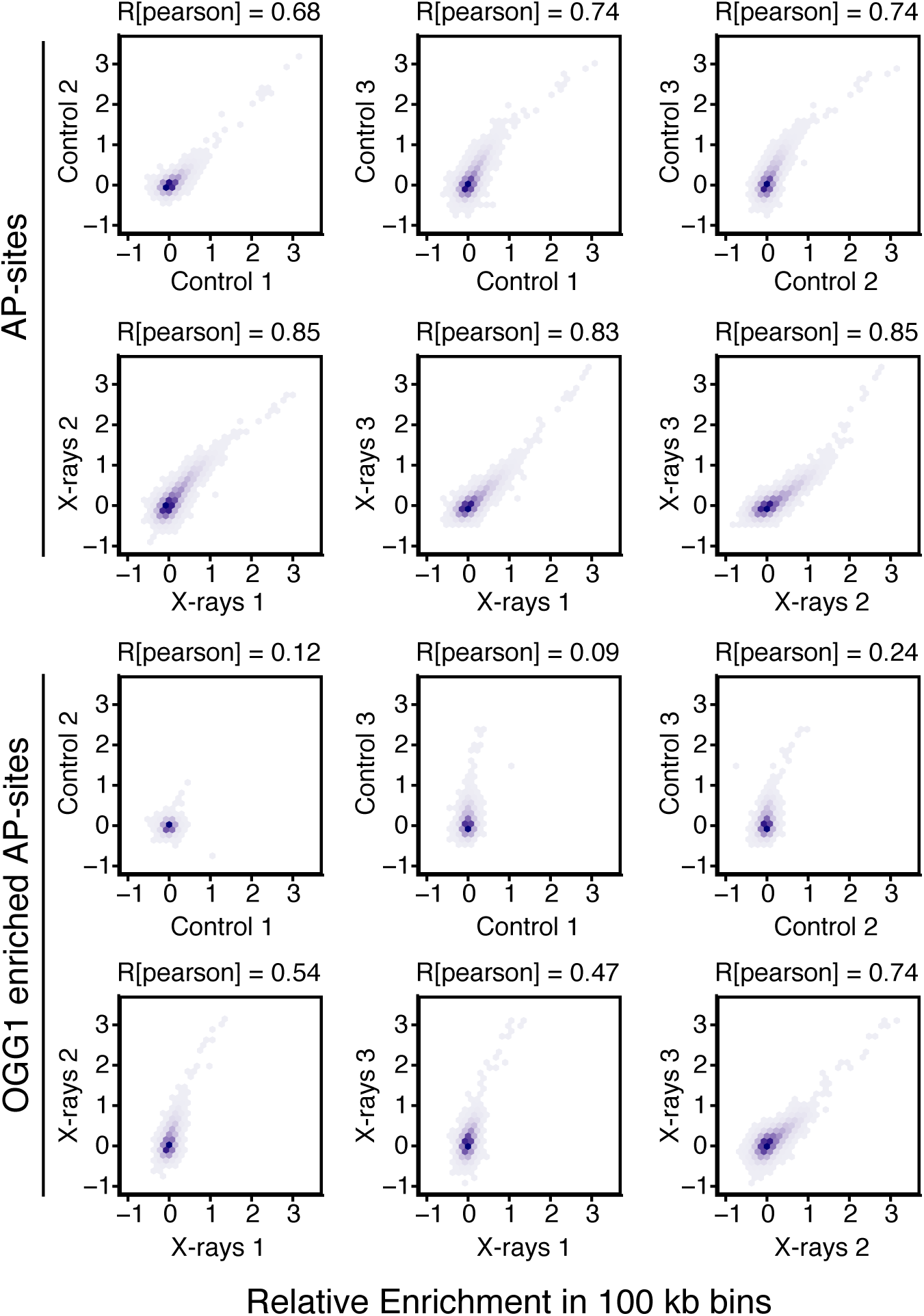
Correlation of treatment replicates at 100kb resolution. Relative enrichment was correlated for each replicate in 100kb resolution using Spearman correlation. Depicted is the relative density of the pairwise correlation for the relative enrichment in all four conditions. Correlation is dependent on non-random distinct distribution patterns. Therefore, the conditions differ in their correlation coefficients dependent on the distinctness of enrichment patterns, with AP-sites and X-ray treatment representing the highest correlation coefficients.

## 8. Methods

### 8.1 Cell culture and X-ray treatment

HepG2 cells were chosen for these experiments on the basis of the availability of additional data from the ENCODE project. In addition, HepG2 cells are preferentially used for DNA damaging compounds that require enzymatic activation (e.g. aflatoxin), which may allow comparison of pathways and damage types in later studies.

HepG2 cells were cultivated at 37°C and 5% CO2 in Dulbecco’s Modified Eagle Medium(DMEM; Invitrogen) supplemented with 1% essential amino acids, 1% pyruvate, 2% penicillin/streptavidin and 10% heat-inactivated fetal bovine serum (FBS). ∼1×10^6^ cells were exposed to 6Gy X-ray using a SOFTEX M-150WE in triplicates. Triplicate samples of untreated control cells were processed in parallel, excluding irradiation. Cells were harvested 30 minutes post-treatment.

Successful treatment was confirmed using immunocytochemical staining for gH2AX. Cells were fixed in 2% formalin in phosphate buffered saline pH 7.2 (PBS). Blocking and permeabilisation were performed with 0.2 % fish skin gelatin, 0.5 % bovine serum albumin (BSA), 0.5 % Triton X-100 in PBS. Staining for γgH2AX was done with a mouse monoclonal antibody (Millipore #05 636) in 1:2000 dilution and stained with a FITC coupled secondary antibody. Nuclear staining with DAPI was included in the mounting medium (ProLong Gold Antifade Mountant, ThermoFisher, Catalogue Number P36931). Images were taken with an Olympus FV1000 microscope.

### 8.2 In vitro pulldown of damaged oligonucleotides

Oligonucleotides with defined damage sites were used to determine the efficiency of the pulldown in vitro. The sequence was adapted from the 59mer used by Guibourt et al ^30^ with additional M13 primer binding sites (Table 1).

**Table 1:**
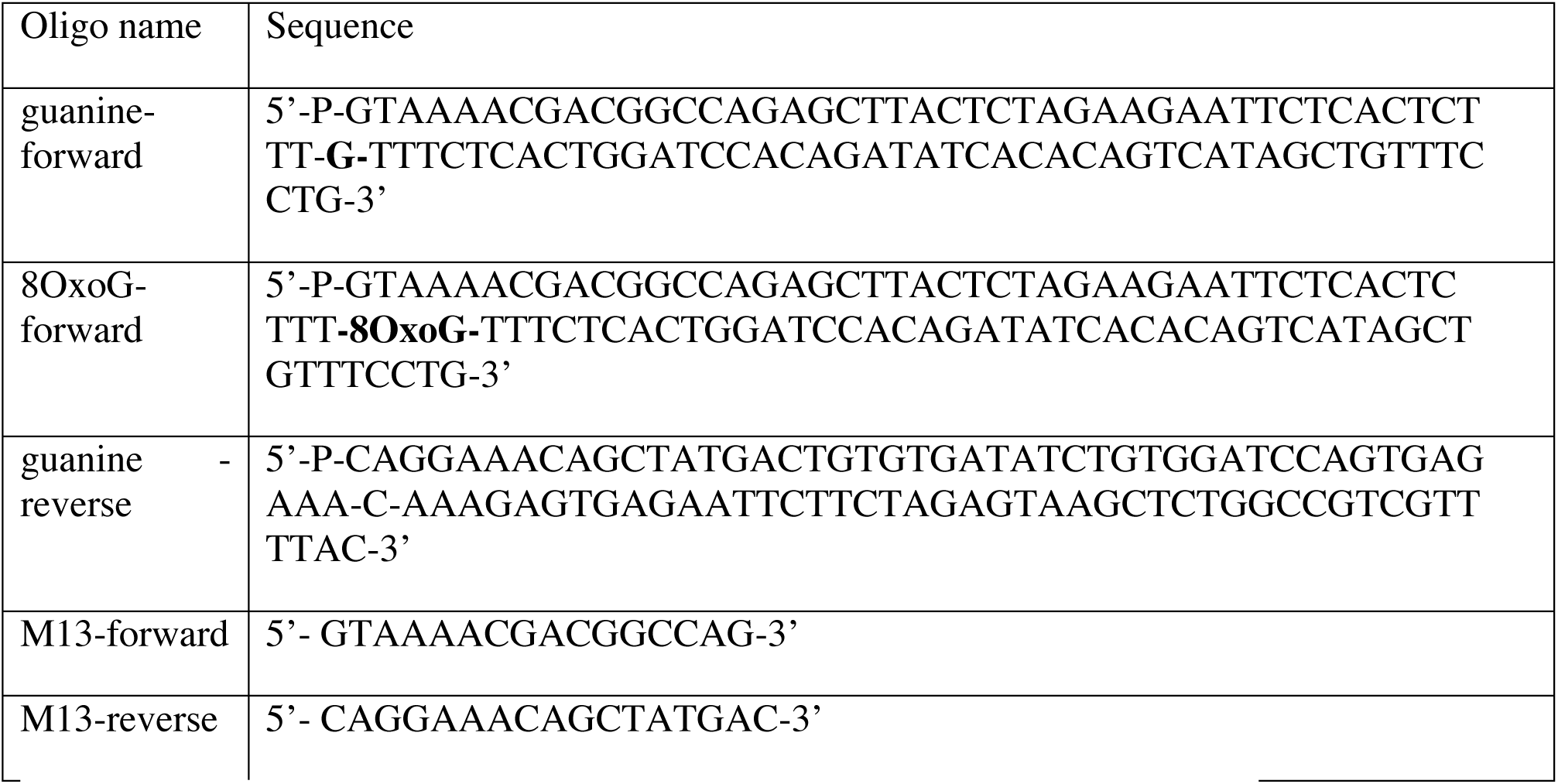
Oligonucleotides and primers used for *in-vitro* pulldown experiments.

Oligonucleotides were hybridized at a concentration of 50 *μ*M for 2 min at 94°C and gradually cooled to room temperature. 10 pmol of the double-stranded 8OxoG-containing oligonucleotide was enzymatically digested with one unit recombinant OGG1 (New England Biolabs, catalogue number M0241L) in New England Biolabs (NEB)-buffer 2 and bovine serum albumin (BSA) and simultaneously tagged with biotin using 5 mM Aldehyde Reactive Probe^31^ (ARP; Life Technologies, catalogue number A10550) for 2h at 37°C. The control oligonucleotide with guanine was tagged with biotin using 5 mM ARP in TE-buffer containing 10mM Tris and 1mM EDTA, pH8. Samples were purified using a ChargeSwitch PCR Clean-up Kit (Invitrogen, catalogue number CS12000).

Half of the sample (up to 5 pmol) was saved as input. The other half was processed for pulldown using 5 μl MyOne Dynabeads (Life Technologies, catalogue number 65601). Beads were washed3x with 1M NaCl in TE-buffer and re-suspended in 2M NaCl in TE-buffer and then added to the equal volume of oligonucleotide solution. The pulldown was performed for 10h at room temperature. The beads were washed 3x with 1M NaCl in TE-buffer. To release the DNA from the beads, the beads were incubated in 95% formamide, 10mM EDTA for 10 min at 65°C and subsequently purified using the ChargeSwitch PCR Clean-up Kit. 2 % of the pulldown was used as template for qPCR. Q-PCR was performed in 25 *μ*l reactions using a Biorad CFX96 Real-Time System with 2x Maxima SYBR Mastermix (ThermoFisher, K0221) and 0.3uM primers. Of the saved input, 1 % was used as template for qPCR.

Recovery of Input was calculated as 2^−ΔCT^ with the differential between pulldown and input. The data were subsequently normalised to the guanine-oligonucleotide as it represents the background pulldown efficiency including background from spontaneous AP-sites that presumably arise as a result of the heating step used to anneal the oligonucleotides.

### 8.3 AP-site colorimetric measurement and AP-Seq

Total genomic DNA was extracted using a Blood and Tissue Kit (Qiagen, catalogue number 69506) and genomic DNA was kept on ice during the process. Antioxidants were not applied in this experiment to avoid artefacts through sequence specific effects. Since treated samples and the untreated control are exposed to the same technical artefacts from sample processing, these should be accounted for in the data analysis. 5.7μg of genomic DNA was tagged with biotin using 5mM Aldehyde Reactive Probe ^31^ (ARP; Life Technologies, catalogue number A10550) in phosphate buffered saline (PBS) for 2h at 37°C. Genomic DNA was then purified using AMPure beads (Agencourt, catalogue number A63882) with 1.8x bead solution, 2x 70% ethanol washing; beads were not allowed to dry to prevent DNA from sticking.

Colorimetric measurement of AP-sites was performed using a commercial Kit (abcam, catalogue number ab65353) following the manufacturers protocol starting from the DNA binding step with 60 μl and 0.1 *μ*g/μl. Optical density at 650 nm was normalized using the standard curve of defined damage sites. From the resulting values, the log2 fold difference to the control mean was calculated and depicted as mean and standard error of the mean. These data were not used for normalisation purposes of the sequencing experiments due to the general semi-quantitive nature of this method.

For AP-Seq, biotinylated DNA was fractionated using a Covaris fractionator in 130μl for a mean fragment length of 300bp. After separating 30μl for sequencing as the input sample, the remaining DNA was used for biotin-streptavidin pulldown, using MyOne Dynabeads (Life Technologies, catalogue number 65601). 120μl beads (10μl per sample) were washed 3x with 1μl 1M NaCl in Tris-EDTA buffer (TE buffer) and re-suspended in 100μl 2M NaCl in TE and then added to 100μl of the sonicated DNA. Samples were rotated at room temperature for 10h. Subsequently the beads were washed 3x with 1M NaCl in TE and finally re-suspended in 50μl TE for library preparation.

For the *in vitro* OGG1-enrichment, 10μg of genomic DNA was digested with recombinant OGG1 (New England Biolabs, catalogue number M0241L). 0.1μg enzyme was taken for 1μg of genomic DNA in New England Biolabs (NEB)-buffer 2 and bovine serum albumin (BSA) for 1h, 37°C. Such conditions for the enzymatic digest should account for sequence content dependent differences in enzyme activity as described by Sassa et al ^35^. Digested DNA was subsequently purified using AMPure beads as described above. The DNA was subsequently tagged with ARP as described above.

### 8.4 Library preparation and sequencing

Both the damage-enriched and input DNA were *in vitro* repaired using PreCR (NEB catalogue number M0309L). The input DNA and supernatant of the pull-down were purified using AMPure beads. The purified pull-down was recombined with the beads and library preparation was performed on the re-pooled sample containing the supernatant and the beads. A 125bp paired-end ChIP-Seq library preparation kit (KAPA Biosystems catalogue number KK8504) was used and sequencing performed using an Illumina HiSeq 2000 on first a rapid and then a high-output run (catalogue number FC-401-4002). The resulting data were subsequently combined.

### 8.5 Read processing library normalisation and damage quantification

Unless stated, data-processing was performed using R 3.4.0 and Bioconductor 3.5. The quality of damage-enriched AP-seq samples (n=12) and corresponding input samples were checked using FastQC (https://www.bioinformatics.babraham.ac.uk/projects/fastqc/); the quality was sufficient that no further filtering was required before alignment. The reads were mapped to the reference human genome (version hg19) using the Bowtie2 algorithm (http://bowtie-bio.sourceforge.net/bowtie2/index.shtml) ^57^ with standard settings, allowing for 2 mismatches and random assignment of non-uniquely mapping reads. To confirm the robustness of key results, analyses were repeated excluding non-uniquely mapped reads (reads with FLAG 3 filtered using SAMtools; http://samtools.sourceforge.net/) ^58^. Data were visualised with the Integrative Genomics Viewer version 2.3.92 (http://software.broadinstitute.org/software/igv/) ^59^.

Paired reads were imported into R using the “GenomicAlignments” and “rtracklayer”^60^ packages. Paired reads mapping more than 1kb apart were discarded. Filters were applied to assess read duplication, reads mapping to the Broad Institute blacklist regions (ftp://encodeftp.cse.ucsc.edu/users/akundaje/rawdata/blacklists/hg19/wgEncodeHg19ConsensusSignalArtifactRegions.bed.gz) ^61^, and whether reads overlap with repeats annotated in the UCSC RepeatMasker track from the UCSC Table Browser (rrmsk_hg19.bed). The main analysis was performed without applying these filters, but the robustness of key results was confirmed by repeating analyses with the filters.

Inter-library normalization was performed using only genomic areas of low damage. It was necessary to consider that increased exposure to DNA damage leads to increased library sizes. A global scaling factor was calculated as the mean read coverage in a low-damage subset (10 %) of 100kb bins, which were identified by their read coverage as the lowest decile of 100kb bins over the mean of all samples.

Relative Enrichment of DNA damage was assessed through the normalised log2 fold-change of the enriched sample over input (termed Relative Enrichment). This should account for biases derived from DNA amounts after genomic DNA extraction, as well as GC content biases from sequencing, which would affect the pull-down samples and inputs alike. Analyses were restricted to Chromosomes 1 to 22 and X, except for the 100kb damage distribution map which includes the Y chromosome (Figure 1B).

All analyses were performed using the average Relative Enrichment in appropriate bin sizes tiled across the genome or covering genomic elements. For a large scale overview a bin size of 100 kb was chosen for comparability with related studies ^19,20^. Genome browser images were generated using absolute read counts pooled over replicates. Peak calling was generally not performed as it was deemed inappropriate for this type of data.

Each treatment condition was independently used for relative comparison within the samples. Lack of absolute quantification suggest that instead of using primary AP-sites as input for OGG1-enriched AP-sites, it is more appropriate to show them side-by-side for comparison.

Correlation of biological replicates was assessed using Pearson correlation in 100kb resolution (Sup. Figure 3).

### 8.6 Analysis on local oxidative damage distribution

The karyogram map was compiled using the mean of the replicates at 100kb resolution with “ggbio” ^62^ karyogram plot fixing the colour scale to a Relative Enrichment of −1 to 1. Enrichment over chromosomes was also depicted with 100kb resolution for the mean of the replicates with shades depicting the standard error of the mean of triplicates. For illustration purposes data were smoothed with a Gaussian smooth over 10 bins, using the smth.gaussian function of the “smoother” package. Correlations at 100kb resolution were performed using Spearman correlation. Fine resolution images were depicted using the IGV browser without any additional smoothing applied.

### 8.7 Epigenome and feature analysis

Genome-wide feature sets were obtained from the UCSC Genome Browser. Chromatin features for HepG2 cells were retrieved from the data repository generated in the context of the ENCODE consortium and obtained through https://www.encodeproject.org/ ^61^. Where applicable, datasets were pooled. Accession numbers are listed in Table 2.

**Table 2:** HepG2 specific datasets obtained from ENCODE and genomic annotation datasets obtained from the UCSC browser

| Feature | ENCODE accession numbers, URL, or UCSC table browser ID |
| --- | --- |
| DNase hypersensitivity | ENCFF774LVT |
| H3K4me3 | ENCFF000BGT |
| H3K4me2 | ENCFF000BFV |
| H3K4me1 | ENCFF000BFC |
| H3K27me3 | ENCFF001FLH, ENCFF001FLI |
| H3K9me3 | ENCFF000BEW |
| H2Az | ENCFF000BEK |
| H4K20me1 | ENCFF000BFJ |
| H3K36me3 | ENCFF001FLR, ENCFF001FLS |
| H3K79me2 | ENCFF000BGB |
| H3K27ac | ENCFF000BGH |
| H3K9ac | ENCFF000BGM |
| RNA-Seq | ENCFF000DPL, ENCFF000DPM, ENCFF000DPN, ENCFF000DPO |
| Replication timing | http://hgdownload.cse.ucsc.edu/goldenpath/hg19/encodeDCC/wgEncodeUwRepliSeq/wgEncodeUwRepliSeqHepg2WaveSignalRep1.bw |
| Mappability | http://hgdownload.cse.ucsc.edu/goldenpath/hg19/database/wgEncodeCr gMapabilityAlign100mer.bigWig |
| Transcription factor binding sites | http://hgdownload.cse.ucsc.edu/goldenpath/hg19/database/tfbsConsSites.txt |
| CTCF binding sites | ENCFF661OYF |
| DNase hypersensitivity sites | wgEncodeAwgDnaseUwdukeHepg2UniPk.bed |

Transcript density was calculated through the genome coverage with any one transcript as defined by UCSC. Distance to telomeres and centromeres was calculated as the absolute base pair distance to annotated telomeres and centromeres.

Genomic and chromatin features were calculated as mean values in 100kb bins over the genome and clustered using hierarchical clustering of Spearman’s correlation coefficients. Features were then correlated (also Spearman) to the individual DNA damage levels. Data points represent the mean of the correlation coefficients with the standard error of the mean over replicates.

### 8.8 GC content analysis

GC content preference of DNA damage distribution was assessed at 1kb resolution. For each 1kb bin in the genome, GC content was calculated and rounded to the closest percentage. Bins with more than 10% undefined sequence were censored. For all bins falling into a particular percentage range, mean Relative Enrichment was calculated with also the standard error of the mean for biological replicates. Averaging over the bins in each category accounts for the lower numbers of bins with extreme GC content. For display, a Gaussian smooth was applied reaching over 10% GC content range.

### 8.9 DNA damage distribution over gene profile

Metaprofiles over coding genes were compiled using the UCSC transcript annotation. The mean was taken for different elements of the genes, which are comprised of a total of 26,860 transcripts. Gene elements were either centred around an appropriate centre point, in which case the mean Relative Enrichment was calculated for each base pair in the respective region. For gene elements of different sizes the mean over the gene element was taken. Independent of their size they were weighted as equal in subsequent analyses. The metaprofile was then compiled with the different gene elements in the following order: 48,838 promoters were centred around the transcriptional start site with 1kb sequence in 5’ direction and 500bp in 3’. 58,073 5’-UTRs, 214,919 exons, and 182,010 introns were addressed as a scaled mean. In addition, exons and introns were addressed through the exon-intron junction, both 5’ and in 3’ of the exon +/-250bp. Given the small sizes of exons, 250bp partially also contains following gene elements. The end of genes is represented through the means of 28,590 3’-UTRs and 43,736 transcription termination sites with 500bp in 5’ direction and 1kb in 3’. 22,480 intergenic regions were addressed as the mean of each region. Shades represent the standard error of the mean over biological replicates.

### 8.10 GC content and transcription dependent promoter analysis

Gene transcription was assessed using RNA-Seq data for HepG2 cells from the ENCODE consortium (Table 2). Replicates were pooled and RNA-Seq coverage was calculated for each unique UCSC defined transcript (n = 57,564). Promoters, i.e. the transcriptional start sites +/-1kb for each transcript were grouped into 11,058 silent promoters and the remaining 46,506 into deciles of increased transcriptional use. In parallel, the mean GC content for each promoter was calculated, which were then also grouped into deciles based on their GC content. Mean damage was assessed for each promoter in these groups.

### 8.11 Retrotransposon analysis

Retrotransposon information was obtained from the UCSC repeat masker. For repetitive sequences, there is a risk of mapping issues and errors of annotation. Therefore, retrotransposon analysis was limited to families of these repeats, where location issues should not arise and mis-estimation of total repeat numbers should largely be balanced out through the IP vs. input comparison. Analyses for particular locations was restricted to the shorter *Alu* repeats, where mapping issues should be minimal and the findings were confirmed by excluding ambiguous mapping.

LINE elements were defined as belonging to *LINE* element families of L1PA7 or newer and only considered, if the size fell between 5.9 and 6.1kb (n=2,533). Alus were considered when 270 to 330bp in size (n=848,350). Retrotransposons were anchored to their start sites and addressed with flanking regions from the start −1kb to +7kb for *LINE* elements and −200bp to +500bp for *Alu* elements. Metaprofiles were compiled as the mean Relative Enrichment over the respective region. GC content was assessed as the mean GC content at the particular site and smoothed using Gaussian smoothing in windows of 5% of feature length.

### 8.12 Transcription factor binding sites, CpG islands and G-quadruplex structure analysis

Transcription factor binding sites were obtained as the consensus set from ENCODE (Table 2), which is cell line unspecific. (n=5,717,225). HepG2 cell specific CTCF binding sites (n=48,671) and DNase hypersensitivity sites (n=192,735) were obtained through ENCODE and UCSC respectively (Table 2). G-quadruplex (G4) structures were obtained using the G4Hunter method ^63^, utilising directly the reference file QP37_hg19_ref.RData provided with the associated R package (n=359,446) with the exception of telomeric G4 structures with the centre less than 500bp away from the chromosome end (n=3). CpG islands were defined through UCSC (n=27,443). Features were considered to be in a promoter, if they overlap with the region of a transcriptional start site +/-1kb. They were considered to overlap with DNase hypersensitivity only when the feature itself overlaps with a DNase hypersensitivity site. For metaprofiles the centres of the features were considered and mean Relative Enrichment of damage levels assessed relative to the centre point. For quantification of mean damage at a given feature site, only the feature itself was addressed and quantified as the mean Relative Enrichment over the region. The GC content of transcription factor binding site was however calculated as the mean over the region (+/-500bp) around the transcription factor binding site. Groups of features were summarised using the median.

### 8.13 Telomere analysis

Due to expected mapping artefacts at telomeric repeats, telomeres were addressed separately not using the aligned sequence. Instead, Telomere hunterversion 1.0.4.(https://www.dkfz.de/en/applied-bioinformatics/telomerehunter/telomerehunter.htμl) ^64^ was used to filter out reads that map to telomeric repeats. These were reassigned to intratelomeric and subtelomeric regions or other locations. Of these, only the intratelomeric repeats were considered. Normalisation between libraries was performed not within the Telomerehunter package but separately with the global scaling factor as described above using only genomic areas of low damage. The global scaling factor was calculated as the mean read coverage in a low-damage subset (10 %) of 100kb bins, which were identified by their read coverage as the lowest decile of 100kb bins over the mean of all samples. Mean Relative Enrichment between biological replicates was calculated with the standard error of the mean.

### 8.14 Microsatellite analysis

Microsatellites were defined through the UCSC repeat masker as the “Simple _repeat” class. For quantification purposes, reverse complement repeat classes were combined. Only microsatellite sequences that are represented >1,000 times in the genome were considered. This leaves 39 repeat types, which are represented by a total of 388,350 repeats. Since the damage assessment does not allow strand specificity, repeats were pooled with their reverse complement assigning both orientations to the alphabetically first repeat. Median Relative Enrichment of damage was quantified over each microsatellite type.

### 8.15 Patient selection for mutation analysis

Data for mutations in cancer were obtained from the Pan-cancer Analysis of Whole Genomes consortium ^50^. Contributions of mutational signatures were provided by PCAWG working group 7.^54^

The data set is comprised of 2,702 tumour-normal pairs for 39 cancer types. From this dataset, we obtained all data on mutation rates and mutation signature contributions, as well as clinical metadata. The analysis was restricted to chromosomes 1 to 22 and X. It was focused on C-to-A mutations as this is the major mutation type derived from oxidative damage. This includes the reverse complement G-to-T, as the analysis is not performed strand specifically. Effects from selection processes were not taken into consideration, as the consequences from the average 2.9 driver SNVs per tumour ^65^ on the mutation patterns should be negligible.

For the mutations dependent on oxidative damage, 8 samples were selected that have a polymerase epsilon proofreading defect as determined by a hypermutator phenotype (C-to-A >100,000) with prominence of Signature 10 confirmed as being linked to coding mutations in Pol E. In total, these samples contain 3,436,531 mutations. For information to individual patients see Table 3.

**Table 3:** Selected tumour samples with polymerase epsilon proofreading defect.

| Sample IDs<br>[tumor_wgs_aliquot_id] | Total C-to-A | Proportion C-to-A | Total mutation count | Cancer type | Proportion Signature 10 |
| --- | --- | --- | --- | --- | --- |
| 00aa769d-622c-433e-8a8a-63fb5c41ea42 | 115,337 | 0.46 | 252,195 | ColoRect-AdenoCA | 0.65 |
| 0980e7fd-051d-45e9-9ca6-2baf073da4e8 | 396,377 | 0.44 | 907,411 | ColoRect-AdenoCA | 0.60 |
| 14c5b81d-da49-4db1-9834-77711c2b1d38 | 989,958 | 0.40 | 2,502,427 | ColoRect-AdenoCA | 0.57 |
| 154f80bd-984c-4792-bb89-20c4da0c08e0 | 126,870 | 0.45 | 280,527 | ColoRect-AdenoCA | 0.63 |
| 2df02f2b-9f1c-4249-b3b4-b03079cd97d9 | 964,307 | 0.38 | 2,570,161 | ColoRect-AdenoCA | 0.41 |
| 6ca5c1bb-275b-4d05-948a-3c6c7d03fab9 | 243,272 | 0.28 | 871,206 | ColoRect-AdenoCA | 0.55 |
| 93ff786e-0165-4b02-8d27-806d422e93fc | 436,686 | 0.43 | 1,024,918 | ColoRect-AdenoCA | 0.50 |
| b0a83df8-dd2c-4c1b-b238-9081d2c22258 | 163,724 | 0.54 | 303,201 | Uterus-AdenoCA | 0.65 |

For comparison, all other 2,695 tumour samples were taken with a total of 6,008,940 C-to-A mutations.

Tumour samples with mutations in direct processing of 8OxoG or AP-sites were identified through assessing, whether mutations fall into the coding sequence of OGG1 (n=7), APEX1 (n=3), or FEN1 (n=3). Mutations were considered, if their effect determined by the ensembl VEP tool (http://www.ensembl.org/Multi/Tools/VEP) ^66^ identified them as missense variants, stop codon gained, frameshift variants, or splice donor variant. They were not considered, if there was an underlying hypermutator phenotype of >100,000 C-to-A mutations (n=2). Information to individual patients can be found in Table 4.

**Table 4:** Selected tumour samples with coding mutations in OGG1, APEX1, of FEN1.

| Sample IDs<br>[tumor_wgs_aliquot_id] | Total<br>C-to-A | Proportion<br>C-to-A | Total<br>mutation<br>count | Cancer<br>type | Coding<br>Mutation |
| --- | --- | --- | --- | --- | --- |
| 42f88b95-fa12-47c7-93f1-cf72f207291c | 1,308 | 0.16 | 7,945 | Kidney-RCC | OGG1 |
| 4a1ad661-f6ae-44e8-b50b-72ff658ff22b | 1,480 | 0.19 | 7,872 | CNS-GBM | OGG1 |
| 7456abd5-303e-4e6f-bf4e-47efefc7310f | 913 | 0.16 | 5,729 | Breast-AdenoCA | OGG1 |
| dc4ba4bc-6333-4fe9-8805-e058cc9e6e18 | 1,963 | 0.17 | 11,509 | Panc-Endocrine | OGG1 |
| e6801359-d1d7-4871-b2fb-180674a2e469 | 1,506 | 0.16 | 9,141 | Kidney-RCC | OGG1 |
| f7e7d61f-e2dc-b523-e040-11ac0c482000 | 477 | 0.16 | 3,011 | Breast-AdenoCA | OGG1 |
| fc5dc6d8-62d2-76d8-e040-11ac0d4863c3 | 1,494 | 0.17 | 8,637 | Breast-AdenoCA | OGG1 |
| 45a7949d-e63f-4956-866c-df51257032de | 2,631 | 0.10 | 25,181 | Bladder-TCC | APEX1 |
| 9ebac79d-8b38-4469-837e-b834725fe6d5 | 2,594 | 0.16 | 16,191 | Panc-AdenoCA | APEX1 |
| bf91afc4-aa2b-4365-80c5-b98c9d118e10 | 333 | 0.13 | 2,543 | Panc-Endocrine | APEX1 |
| 369c06f2-8904-49cb-99d1-dd297ed0cd0c | 3,424 | 0.17 | 20,565 | Lung-SCC | FEN1 |
| 81b1e78c-6032-4ff4-b52a-83456b9450ea | 9,708 | 0.33 | 29,682 | ColoRect-AdenoCA | FEN1 |
| f7b84c09-15d4-3046-e040-11ac0c4847ff | 455 | 0.18 | 2,528 | Breast-AdenoCA | FEN1 |

Patients with oxidative damage induced mutations beyond polymerase epsilon proofreading defects were separated based on the proportion contribution of Signature 18 to C-to-A mutations. Patients were censored that have a hypermutator phenotype (C-to-A >100,000, which includes Pol E proofreading deficient tumour samples) or coding mutations in 8OxoG or AP-site processing as identified above. In addition, patients were also censored based on documented smoking history or previous exposure to chemotherapy/radiotherapy. A total of 2,401 samples were used for analysis. They were grouped into Signature 18 based groups of <10% (n=1,398), 10% to 40% (n=322), 40% to 60% (n=540), and >60% (n=141).

### 8.16 GC content preferences of mutation rates

For each 1kb bin in the genome, GC content was calculated and rounded to the closest percentage. Bins with more than 10% undefined sequence were censored. Mutations falling into bins of 50% GC content or higher was calculated as proportion of the total C-to-A mutation counts. Assuming equal distribution dependent only on base content, a total of 9% of C-to-A mutations would be expected to fall into such high GC content areas of the genome. The cut-off was determined based on the observed vs. expected ratio for C-to-A mutation rates in each GC content bin. For Pol E proofreading defective tumours this ratio drops below 1 at ∼ 40 %, so a 50 % cut-off was chosen. Drawing the cut-off at 60 % GC content gives equivalent results.

### 8.17 Genomic features analysis

Metaprofiles over genomic features were calculated for the features with the same selection strategy as described above. For this, C-to-A mutations were pooled for each patient group. Mean relative mutation rates over features were calculated as relative C-to-A mutation density normalized to 1,000,000 C-to-A mutations per patient group. The mean over the features was normalized for sequence content of the particular location by dividing with a factor of the local GC content divided by the average of 41%. For display purposes, data were smoothed using a Gaussian smooth spreading over 100bp for the gene body profile, Alus, protein binding sites, CpG islands, and G4 structures. *LINE* elements were smoothed using Gaussian smoothing over 200bp to account for the increased noise originating from the lower frequency of this particular feature.

## 9. Acknowledgements

We are most grateful to Peter Van Loo, Kerstin Haase, Clemency Jolly, Jonas Demeulenmeester, and Maxime Tarabichi for their advice and technical assistance for analysing cancer genomics data. For technical assistance with experiments and sequencing we would like to thank Hiroki Goto, Shinichi Yamasaki, Keigo Hikishima, Rehab Abdelhamid, Panagiotis Kotsantis, Valerie Borel, Graeme Hewitt, Lorea Blazquez, and Mary Bronks. For experimental advice, we would like to thank Aswin Mangerich. For advice on data analysis we would like to thank the Crick Bioinformatics Science Technology Platform, in particular Harshil Patel. This work was supported by the Francis Crick Institute which receives its core funding from Cancer Research UK (FC001110), the UK Medical Research Council (FC001110), and the Wellcome Trust (FC001110). NML is a Winton Group Leader in recognition of the Winton Charitable Foundation’s support towards the establishment of the Francis Crick Institute. NML is additionally funded by a Wellcome Trust Joint Investigator Award (103760/Z/14/Z, the MRC eMedLab Medical Bioinformatics Infrastructure Award (MR/L016311/1). ARP and NML were also supported by funding from the Okinawa Institute of Science & Technology Graduate University and a Wellcome Trust Investigator Award. ARP was funded by a postdoctoral fellowship from the Peter and Traudl Engelhorn Foundation. We thank Peter Van Loo and Jernej Ule for helpful advice and discussions throughout the project.

